# Conditionally essential genes for survival during starvation in *Enterococcus faecium* E745

**DOI:** 10.1101/2020.04.10.036137

**Authors:** Vincent de Maat, Sergio Arredondo-Alonso, Rob J.L. Willems, Willem van Schaik

## Abstract

The nosocomial pathogen *Enterococcus faecium* can survive for prolonged periods of time on surfaces in the absence of nutrients. This trait is thought to contribute to the ability of *E. faecium* to spread among patients in hospitals. Because there is currently a lack of data on the mechanisms that are responsible for the ability of *E. faecium* to survive in the absence of nutrients, we performed a high-throughput transposon mutant library screening (Tn-seq) to identify genes that have a role in long-term survival during incubation in phosphate-buffered saline (PBS) at 20°C. A total of 18 genes were identified by Tn-seq to contribute to survival in PBS, with functions associated with the general stress response, DNA repair, metabolism, and membrane homeostasis. The gene which was quantitatively most important for survival in PBS was *usp* (locus tag: EfmE745_02439), which is predicted to encode a 17.4 kDa universal stress protein. After generating a targeted deletion mutant in *usp*, we were able to confirm that *usp* significantly contributes to survival in PBS and this defect was restored by *in trans* complementation. As *usp* is present in 99% of 1644 *E. faecium* genomes that span the diversity of the species, we postulate that this gene is a key determinant for the remarkable environmental robustness of *E. faecium*. Further mechanistic studies into *usp* and other genes identified in this study may shed further light on the mechanisms by which *E. faecium* can survive in the absence of nutrients for prolonged periods of time.

## Introduction

*Enterococcus faecium* is a commensal of the human gut, but has emerged over the last few decades as an opportunistic pathogen which causes infections in hospitalized patients. *E. faecium* infections are often difficult to treat due to the high prevalence of resistance to antibiotics, including virtually all cephalosporins, aminoglycosides, clindamycin, and trimethoprim-sulfamethoxazole [1]. Additionally, resistance to the glycopeptide vancomycin is increasingly widespread in *E. faecium* strains, further complicating the treatment of infections. While the accumulation of antibiotic resistance determinants is a major contributor to *E. faecium*’s emergence as an important nosocomial pathogen, other adaptations, like the ability to form biofilms [2–4] and interact with host extracellular matrix and serum components [5], are also widespread in clinical isolates. *E. faecium* also has the ability to persist for long periods of time on synthetic surfaces like table tops, handrails, doorknobs and other medical surfaces [6–8].The ability of *E. faecium* to spread via fomites is thus proposed to play a critical role in the inter-patient spread of *E. faecium* in hospital settings [9–11]. Many nosocomial pathogens are thought to spread via environmental contamination, but *E. faecium* can survive 3- to 5-times longer on inanimate objects compared to other Gram-positive nosocomial pathogens, such as *Enterococcus faecalis*, *Staphylococcus aureus*, and streptococci [12–14].

In natural environments, bacteria most often exist in a physiological state that is similar to the stationary phase of growth and are adapted to survive harsh conditions [15, 16]. While survival strategies like sporulation are limited to a subset of bacterial species, most bacteria possess the ability to survive in the complete absence of nutrients (e.g. in water or buffers) for weeks, months and in some cases even years [17, 18]. Research into the mechanisms by which bacteria manage to survive these nutrient-limited conditions is currently relatively scarce. Most long-term survival-related studies have been performed in *E. coli* and have only recently have been broadened to other bacteria. Therefore current knowledge extends to only a handful of genes and molecular systems that enhance either survival during or, recovery from prolonged stationary phase [19, 20]. One well described regulation to stress is the stringent response [19–21], which is involved in nutrient deprivation and several other stresses. The stringent response revolves around the synthesis and degradation of the small signal molecules guanosine 5’-diphosphate 3’-diphosphate (ppGpp) and guanosine 5’-triphosphate 3’-diphosphate (pppGpp), collectively named (p)ppGpp. The production of (p)ppGpp is mediated by RelA and SpoT (in *E. coli*) or Rel/Spo homologues (RSH) in other bacteria [19, 22, 23]. In *E. faecalis* two RSH homologues are known, RelA and RelQ, in which RelA is responsible for mounting the main stringent stress response while RelQ maintains base levels of (p)ppGpp during exponential growth [24–27]. (p)ppGpp downregulates transcription of genes involved in growth and division while upregulating genes involved in stress response. RSH genes are conserved among most prokaryotes and are in general the primary enzymes involved in (p)ppGpp accumulation during starvation [19, 23]. Most other stress responses upon nutrient deprivation involve the expression or activity of DNA repair systems, transcriptional regulation, metabolic pathways, and the biogenesis of cell walls and membranes [19, 28, 29].

While survival in the absence of nutrients appears to be an important factor in the spread of *E. faecium* as a nosocomial pathogen, we currently lack a complete picture of the genes involved in these processes. For this study, we have elucidated the *E. faecium* genes involved in survival upon starvation by high-throughput screening of a *mariner* transposon mutant library (Tn-seq) of a clinical *E. faecium* strain upon incubation in a buffer without carbon- or nitrogen-sources at room temperature for seven days. With this study, we generated the first insights into the mechanisms by which *E. faecium* can survive in the absence of nutrients.

## Results

### Effect of nutrients and temperature during prolonged incubation

The vancomycin-resistant strain *E. faecium* E745 was incubated at either 37☐ or 20☐ in the rich medium BHI or in PBS for up to 10 days **(Figure 1)**. By determining viable counts, we showed that cells in BHI were able to survive for a prolonged period of time. Indeed, the stationary phase cultures incubated in BHI at 20°C, showed no reduction in viable counts during the course of the experiments, while at 37°C the cultures were only losing their viability (0.9-log_10_ reduction) after more than 7 days. During incubation in the nutrient-free buffer PBS, significant reductions in viable counts were observed after 2 days of incubation at 37°C and after 6 days at 20°C, but a sub-population of cells remained viable until the end of the experiment.

**Figure 1:**
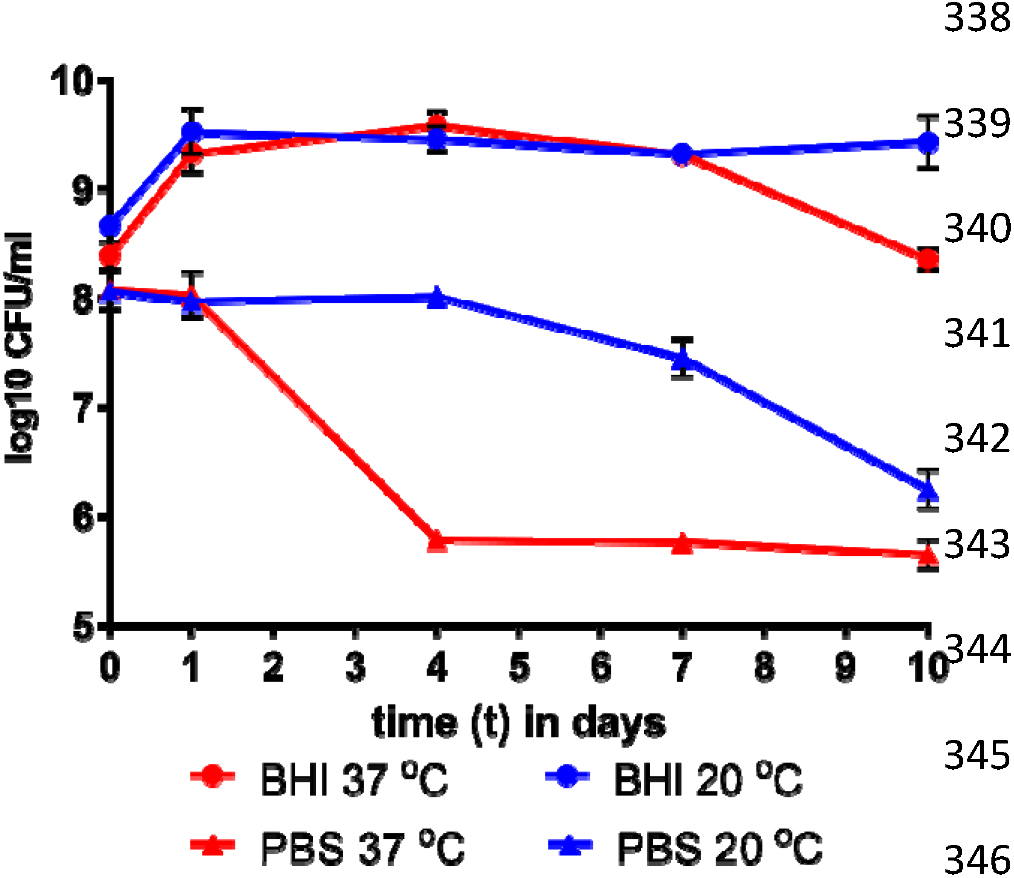
Survival of *E. faecium* E745 during starvation. *E. faecium* strain E745 was incubated for up to 10 days in BHI (circles) or PBS (triangles) at either 37°C (red) or 20°C (blue).

### *E. faecium* E745 genes required for survival during starvation

To investigate which genes were required for survival in PBS at 20☐, we incubated a transposon mutant library of E745 in PBS for 7 days at 20☐. The relative abundance of transposon mutants in 1631 chromosomal and plasmid genes of *E. faecium* E745 at day 7 was compared to the abundance of mutants present at the start of the experiment **(Figure 2)**. The abundance of transposon mutants in 21 genes was significantly reduced (Benjamini-Hochberg corrected P-value of <0.05) by 10-fold or more, indicating that these genes potentially contribute to the survival of *E. faecium* in PBS **(Table 1)**. The predicted functions of these 18 genes consist of 4 main groups, i.e. general stress response, DNA repair, metabolism, and membrane homeostasis. Transposon mutants in four genes had a greatly reduced (more than a 1000-fold) abundance upon 7-day incubation in PBS, indicating that these genes have a large impact on survival in nutrient-limited conditions.

**Figure 2:**
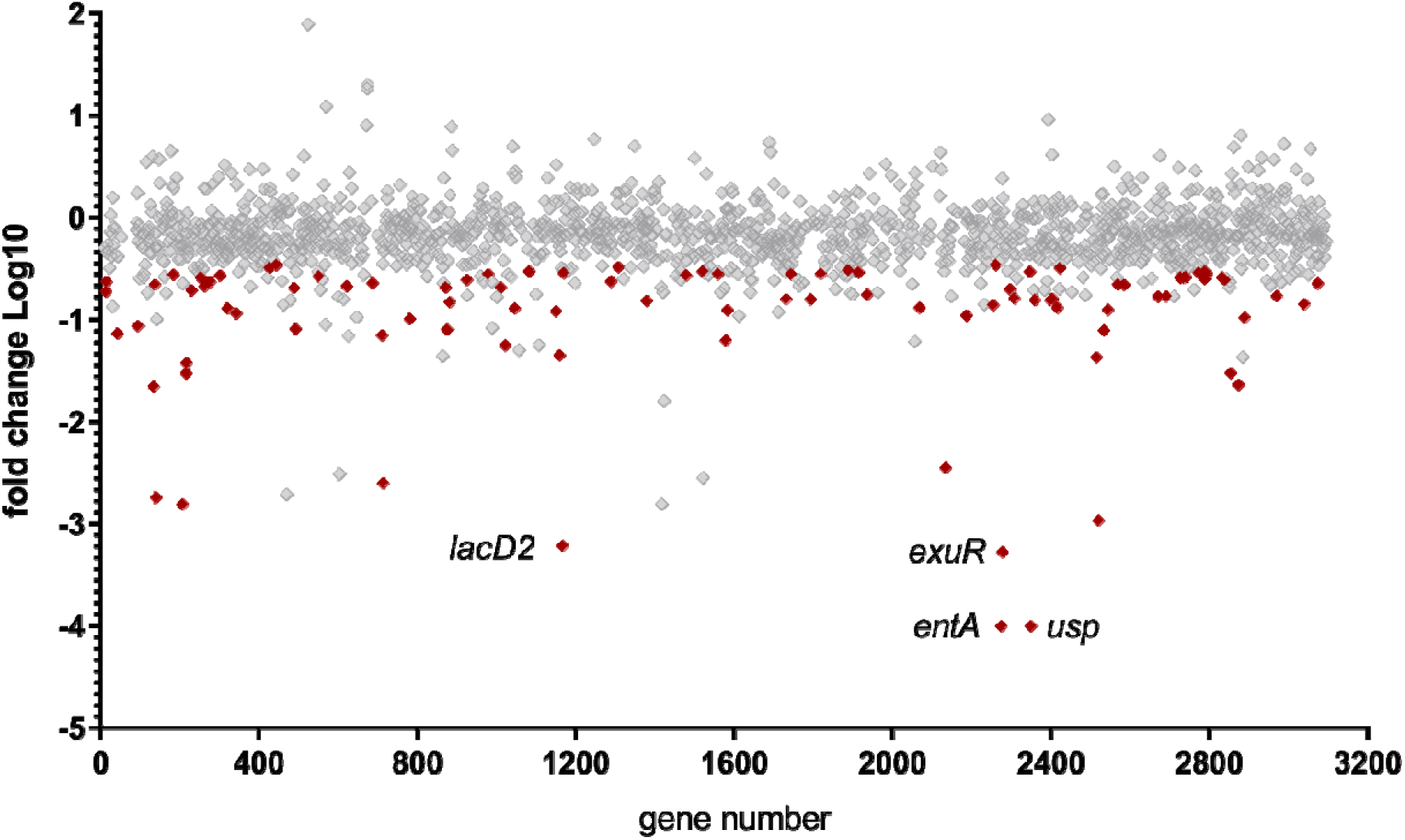
Tn-seq analysis to identify genes with a role in survival during starvation. Each diamond represents a gene represented in the transposon library. The Y-axis indicates the relative abundance of each gene after incubation in PBS at 20°C for 7 days compared to the abundance of each gene at the start of the experiment. A positive value indicates an enrichment of mutants during incubation in PBS while a negative value indicates genes that contribute to survival in PBS, with red diamonds representing genes that passed the threshold for statistical significance. The highlighted dots represent genes that are discussed in the text.

**Table 1:**
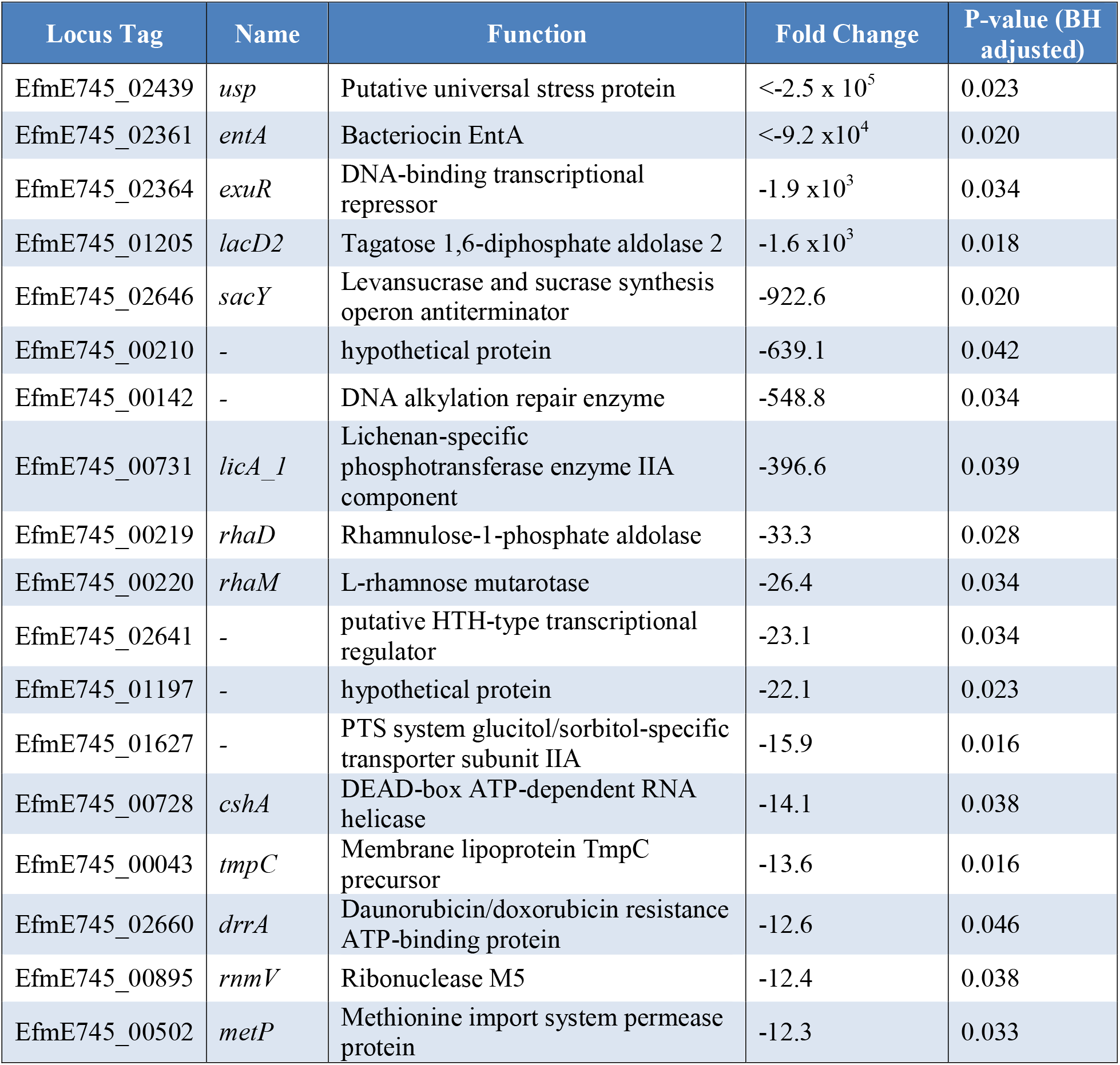
Genes with a role in survival during starvation as determined by Tn-seq. Genes of which the transposon mutants were reduced by more than −10-fold upon incubation in PBS for 7 days, using a Benjamini-Hochberg-adjusted P-value of <0.05 as cut-off for statistical significance.

The *usp* gene (locus-tag: EfmE745_02439) was identified as the most important gene involved in survival in PBS. Inspection of the *mariner* transposon insertion sites in *<u>usp</u>* revealed no unusual patterns in the abundance or placement of the transposon insertion sites, supporting the validity of this hit (**Figure 3A**). The *usp* gene encodes a 17.4 kDa putative universal stress protein with no clear function, although homologs are widespread among all domains of life [30, 31]. In *E. coli*, related proteins are reported to be produced in response to a wide variety of environmental stresses and are highly expressed in growth-arrested cells [30]. The exact mechanism by which these proteins contribute to survival during growth arrest is currently unknown. The second-highest hit in the Tn-seq analyses was found to be in the gene that encodes the previously characterised enterococcal bacteriocin EntA (locus tag: EfmE745_02361) [32, 33]. Like for *usp*, transposon insertions in *entA* were no longer detected in the library at day 7. Bacteriocins are small secreted proteins or peptides produced by bacteria to reduce competition of unrelated bacterial strains. A unique feature of bacteriocins is that they are always co-expressed with their associated immunity gene to avoid auto-inhibition [34]. When we inspected the distribution of transposon insertions in *entA*, we noted that there was only one insertion site in *entA* which is located close to the 3’ end of the gene (**Figure 3B**). As the presumptive bacteriocin immunity gene was found to be overlapping with *entA* and was also disrupted by the transposon insertion, there is a distinct possibility that this transposon mutant is killed due to the production of the EntA bacteriocin during nutrient deprivation, by the other mutants in the population in which these genes were not disrupted. For this reason, we did not included *entA* in further analyses. The transposon mutants in EfmE745_02364 (*exuR*) and EfmE745_01205 (*lacD2*) (**Figure 3C** and **3D**) were also highly reduced upon incubation in PBS, and transposon insertion sites in these genes showed no unusual patterns.

**Figure 3:**
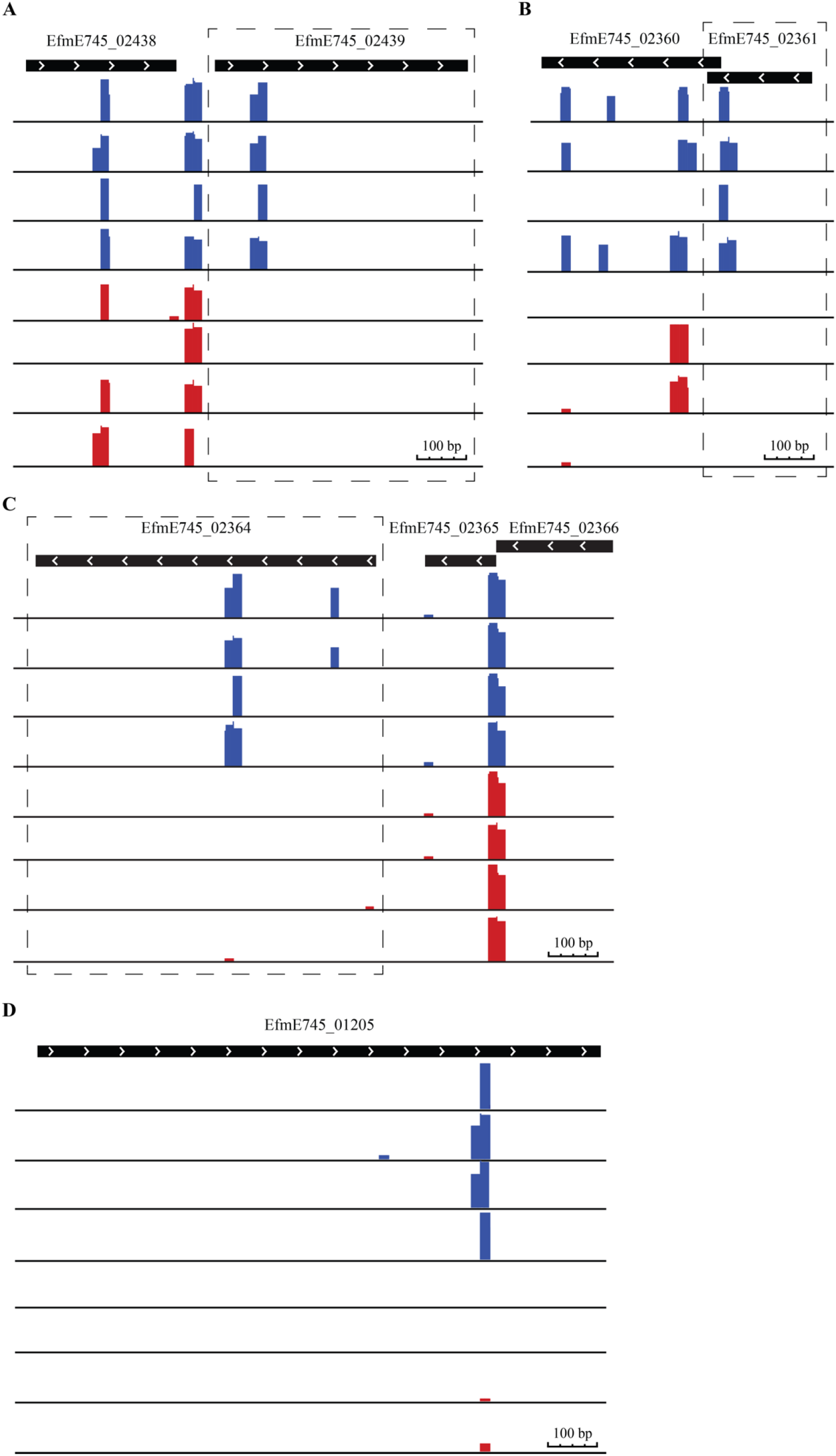
Visual representation of the transposon insertion sites in *usp*, *entA*, *exuR* and *LacD2*. Genes and gene direction are depicted by the black bars with arrows. Transposon insertion abundance is shown by the bars below the genes on a Log_10_ scale. Blue and red bars denote the abundance of transposon insertion mutants at day 0 and at day 7, respectively. Abundance of transposon insertion are shown for EfmE745_02439 (*usp*), EfmE745_02361 (*entA*), EfmE745_02364 (*exuR*) and EfmE745_01205 (*lacD2*) in panel A, B, C and panel D, respectively.

Next we studied the presence of all 18 genes identified by Tn-seq in the whole genome sequences of a collection of 1646 *E. faecium* strains isolated from healthy humans, patients, pets, pigs, and poultry **(Figure 4)**. We found that *usp*, *entA,* and *exuR* are present in 99%, 90% and 100% respectively of the isolates tested and can be thus considered part of *E. faecium* core genome. The *lacD2* gene was only present in 53% of isolates.

**Figure 4:**
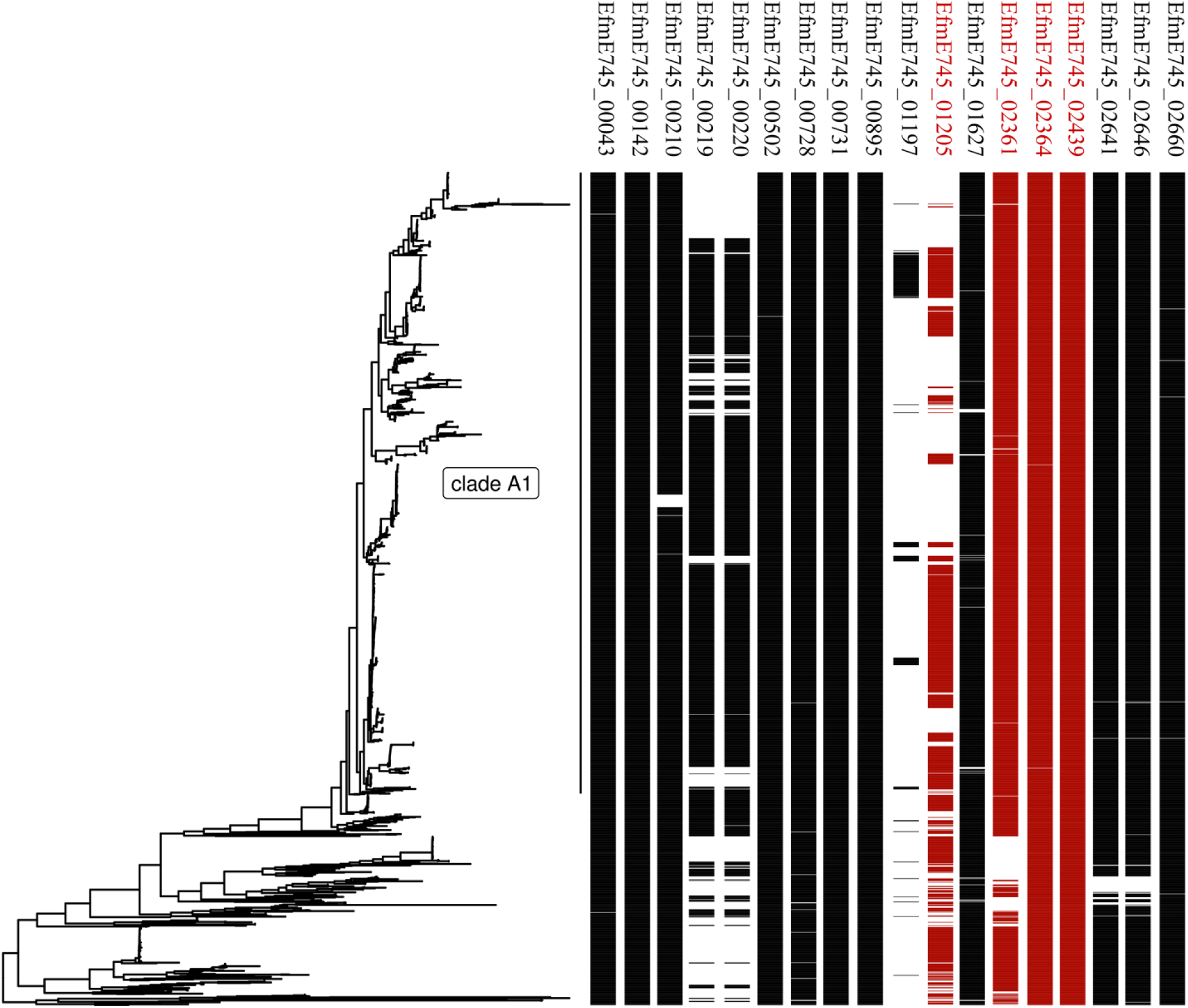
Presence of genes involved in long-term survival under nutrient-limiting conditions in *E. faecium*. Whole genome sequence based phylogenetic tree of 1644 *E. faecium* strains representing the global *E. faecium* population. The presence of the different E745 genes was plotted along the phylogeny using the R package ggtree. Highlighted in red are EfmE745_01205 (*lacD2*), EfmE745_02361 (*entA*), EfmE745_02364 (*exuR*) and EfmE745_02439 (*usp*).

To further test the importance of these genes we have attempted to create gene deletion mutants in EfmE745_02439 (*usp*), EfmE745_02364 (*exuR*) and EfmE745_01205 (*lacD2*), but only managed to construct deletions in *usp* and *exuR*. These mutants were tested for their ability to survive in PBS for a 7-day period.

### The universal stress protein Usp, but not ExuR, contributes to survival of *E. faecium* during prolonged starvation

To determine whether *usp* has an impact on the survival of E745 we created the deletion mutant E745-*Δusp* and complemented the deletion by supplying the *usp* gene *in trans* on a plasmid (strain E745-*Δusp*::*usp*-C). We incubated E745, E745-*Δusp,* E745-*Δusp*::*usp*-C, and E745-*Δusp*::pEF25 (the *usp* deletion mutant transformed with the empty vector used for *in trans* complementation) in PBS at 20°C and determined viability of the cell suspension over a 7-day period **(Figure 5A)**. The *E. faecium* E745 strain showed a similar survival response as seen in **Figure 1.** However, the E745-*Δusp* and E745-*Δusp*::pEF25 cell suspension lost their viability quicker than E745, with a statistically significant (P<0.05; one way ANOVA) 1-log lower viable count after 7 days than the wild-type. The complemented strain, E745-*Δusp*::*usp*-C, responded similar to WT E745 with no statistically significant difference detected at any time point. These results confirm the role of *usp* in survival of *E. faecium* E745 in the absence of nutrients. E745-*ΔexuR* was found to survive equally well as the wild-type strain during starvation (**Figure 5B)**, suggesting that *exuR* does not have a role in survival during starvation.

**Figure 5:**
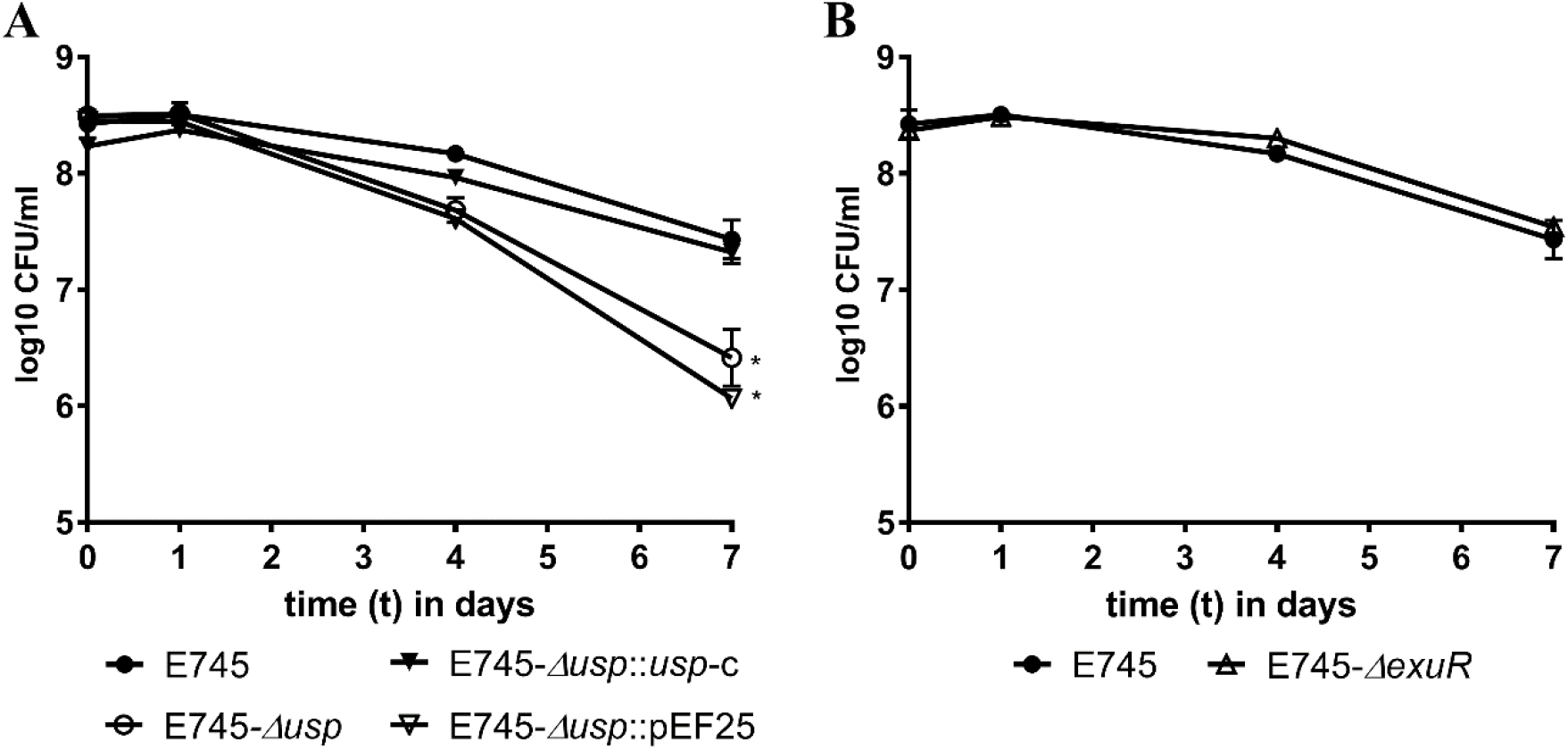
Phenotypic analysis of *usp* and *exuR* deletion mutants. The data represent the data of three biologically independent experiments, with error bars showing standard deviation. All strains were incubated in PBS at 20°C for up to 7 days. * indicates a statistical difference (P<0.05) between survival of *E. faecium* E745 and *Δusp* and *Δusp*::pEF25 at day 7, as determined by one way ANOVA.

## Discussion

In this study we investigated the potential of vancomycin-resistant *Enterococcus faecium* to survive in the absence of nutrients, a condition that *E. faecium* will find itself in when it finds itself on inanimate objects in hospitals. To obtain a better understanding into the genes that are involved in the response upon prolonged starvation, we performed Tn-seq on a transposon mutant library that was starved for nutrients for 7 days at 20°C. This analysis revealed a variety of genes that were potentially involved in retaining viability. Genes could be assigned to four major functional groups, i.e. DNA repair, metabolism, cell membrane and cell wall homeostasis. DNA repair, or lack thereof, can contribute to survival during starvation as reactive oxygen species and other DNA damaging molecules accumulate [35]. Metabolic adaptations during starvation are essential as shifts in metabolism, including changes in transport systems, are required for optimal acquisition of nutrients and the excretion of metabolic waste products [23, 36]. Changes to the cytoplasmic membrane and cell wall have also been described to contribute to the ability of cells to withstand harsh conditions, including starvation [37]. The final category of genes we identified in our Tn-seq analysis have roles in the general stress response, i.e. the cell’s extended toolbox for responding to adverse conditions [30, 31].

We studied the role of the general stress protein Usp (locus tag: EfmE745_0243) which belongs to this last functional group and confirmed that it has a significant impact on survival of *E. faecium* during starvation. Universal stress proteins (USPs) are mostly studied in *E. coli* where they protect the cell against adverse conditions. However, the exact mechanism(s) by which they act are currently unknown. This could be a result of the many different subtypes found, and the complex responses they contribute to. The production of USPs is upregulated during the stringent response in both Gram-negative[38–40] and Gram-positive bacteria [25, 41].

USPs are found throughout all branches of the tree of life and are important for survival under adverse conditions in many species [30, 31]. In this study we expand these observations to *E. faecium*.

The *exuR* gene (locus-tag: EfmE745_02364) appeared to be important for survival upon nutrient starvation on the basis of our Tn-seq screening, but we were unable to confirm this finding by studying an *exuR* deletion mutant. The *E. coli* ExuR homologue represses genes involved in the metabolism of D-galacturonate and D-glucuronate [42, 43]. ExuR expression is upregulated in the absences of glucose which allows *E. coli* to use different carbon sources for its growth and survival. We speculate that ExuR in *E. faecium* is involved in rerouting carbohydrate metabolism, similar to *E. coli*. It is possible that the derepression of pathways involved in alternative carbon source utilization in the *ΔexuR* mutant might negatively affect the growth of cells during the recovery step in the glucose-rich medium BHI, which we performed to minimize a Tn-seq signal originating from dead cells in the cell suspension.

Perhaps surprisingly, we did not identify *relA* or *relQ* in our Tn-seq experiments. We are not aware of studies characterising a deletion, or loss-of-function, mutant in *E. faecium relA* or *relQ* but in *E. faecalis* the *relA* mutant was unable to induce stringent stress response via (p)ppGpp while a *relQ* mutant was still able to induce the stress response but had elevate base levels of (p)ppGpp [16, 27]. Both the *relA* and *relQ* genes of *E. faecium* (locus-tag: EfmE745_02431 and EfmE745_02408) encode proteins with a 83% amino acid identity with RelA and RelQ of *E. faecalis* V583, respectively [24, 26, 27]. In our Tn-seq experiment, we only detected ≤4 transposon insertions in *relA* and ≤ 32 in *relQ* in the different replicates (**Figure S1)**. This low number of transposon insertions suggests that both *relA* and *relQ* contribute importantly to fitness during normal growth and were therefore lost during construction of the transposon mutant library. At this point, we can thus not draw a conclusion on the contribution of *relA* or *relQ* in starvation survival of *E. faecium*.

Most bacteria have a ‘feast and famine’ lifestyle where long periods of severe nutrient limitation are interrupted by short bursts of nutrient availability [28]. For *E. faecium*, we propose that this cycle is particularly relevant for the spread of this opportunistic pathogen in hospitals, where it can survive on inanimate objects until it is transferred to a human host. Indeed, recent work has suggested that this is a trait that characterizes the genus *Enterococcus* and has been selected for over hundreds of millions of years of microbial evolution [37]. The finding that the genes most likely to contribute to the survival of *E. faecium* during starvation are part of the core genome of this species, support the crucial role of starvation tolerance in the evolution of *E. faecium*. While *E. faecium* has a low intrinsic virulence, it has become a significant threat to immunocompromised patients in hospitals due to its ability to rapidly acquire genes involved in antibiotic resistance and other traits that contribute to its ability to successfully colonize patients [44]. The mechanisms by which it can survive and spread in the hospital environment have so far received less attention. In this study, we have identified 18 *E. faecium* genes that could potentially affect its survival in the absence of nutrients. We have confirmed the role of the *usp* gene, but future functional characterization of other genes identified here may further increase our understanding of the mechanisms by *E. faecium* can survive outside human or animal hosts.

## Materials and Methods

### Bacterial strains, plasmids, growth conditions, and oligonucleotides

The vancomycin-resistant *E. faecium* strain E745 [45] was used throughout this study. This strain was isolated from a rectal swab of a hospitalized patient, during routine surveillance of a VRE outbreak in a Dutch hospital, and its genome was previously sequenced to completion [45]. Unless otherwise mentioned, *E. faecium* was grown in brain heart infusion broth (BHI; Oxoid) at 37°C. The *E. coli* strains EC1000 [46] and DH5α were grown in Luria-Bertani (LB) medium. When necessary, antibiotics were used at the following concentrations: erythromycin 50 μg ml^−1^, chloramphenicol 10 μg ml^−1^, spectinomycin 200 μg ml^−1^ and gentamycin 300 μg ml^−1^ for *E. faecium* and spectinomycin 100 μg ml^−1^, chloramphenicol 4 μg ml^−1^, erythromycin 100 μg ml^−1^ and gentamicin 50 μg ml^−1^ for *E. coli*. The transposon library in *E. faecium* E745 was previously described [45]. The vectors pWS3 [47], pVDM1001 [48], pCRE-Lox [49], pEF25 [50] and pGPA1 [45] were obtained from our laboratory’s culture collection. Genomic DNA isolation was performed using the Wizard Genomic DNA Purification kit (Promega).

The sequences of all oligonucleotides used in this study are provided in Table 2. Synthesized DNA fragments are listed in Supplementary Table S1.

**Table 2:**
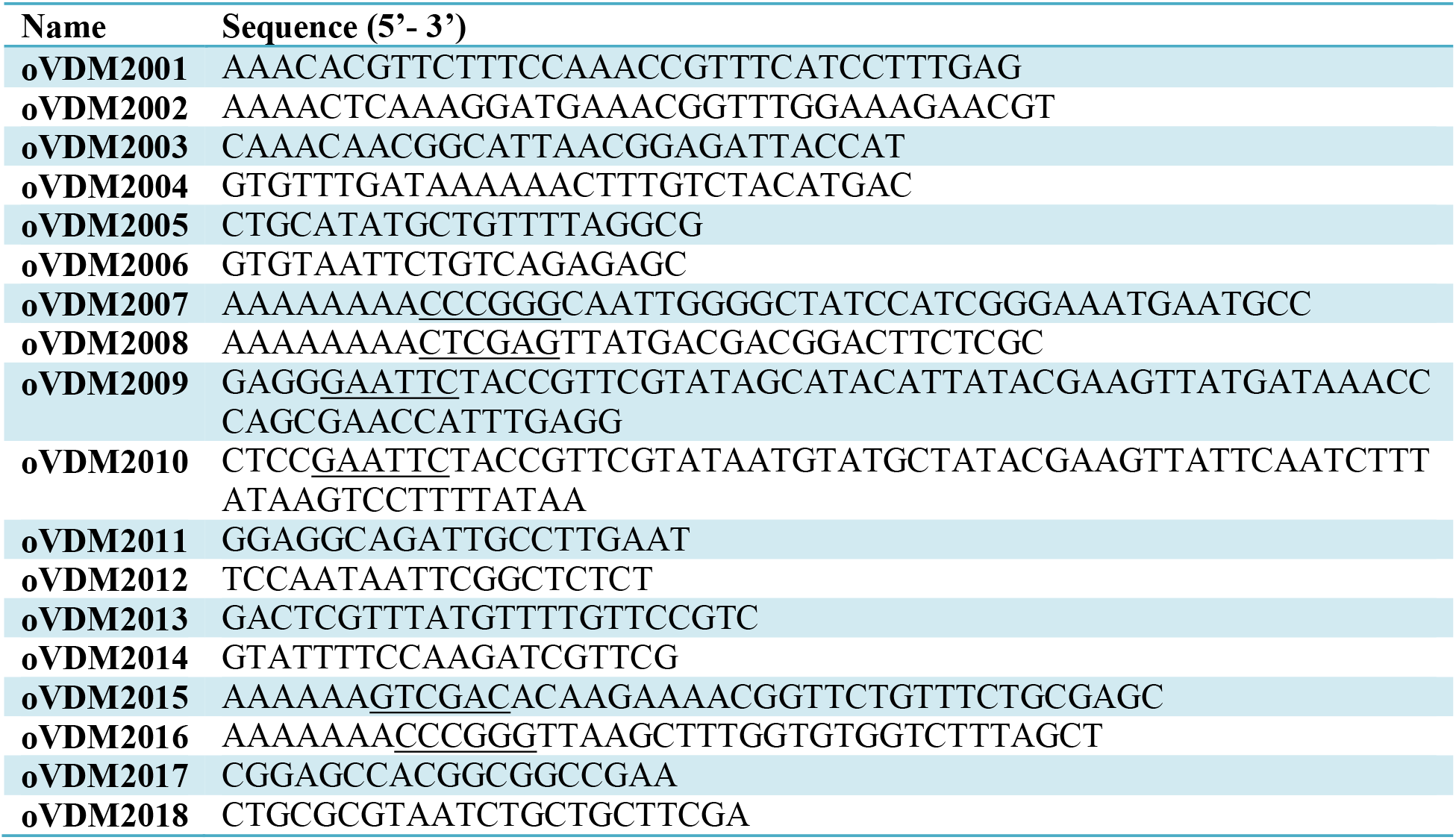
Oligonucleotides used in this study. Relevant restriction sites are underlined.

### Isolation and transformation of plasmids

Plasmid isolation from overnight *E. coli* cultures was performed using the GeneJET plasmid miniprep kit (Thermo Fischer Scientific, Bleiswijk, The Netherlands) according to the manufacturer’s instructions. Transformation of plasmids into *E. faecium* E745 was performed as previously described [49].

### Determination of viability during starvation

*E. faecium* E745 cultures were grown overnight in 3 ml BHI at 37☐ with shaking at 150 rpm. An aliquot of 20 μl of an overnight culture was transferred to 20 ml BHI which was again grown at 37☐ with shaking at 150 rpm until the culture reached an optical density at 600 nm (OD_600_) of 0.3 - 0.4. Aliquots (3 ml) of these cultures were then centrifuged (3000 g, 5 min), washed once in phosphate-buffered saline (PBS; NaCl 137 mM; 2.7 mM KCl; 10 mM Na_2_HPO_4_; 2 mM KH_2_PO_4_; pH 7.4) and resuspended in 3 ml PBS and incubated at either 37☐ or 20☐ with continuous shaking at 150 rpm. Controls remained in BHI and were incubated at the same temperatures with shaking. At days 0, 1, 4, 7, and 10 samples were taken and serially diluted in PBS, after which viable counts were determined using the Miles and Misra method [51].

### Tn-seq analysis of conditionally essential genes in *E. faecium* E745 during prolonged starvation

For the identification of genes that were conditionally essential for prolonged starvation in *E. faecium*, we grew the previously constructed *mariner* transposon mutant library in *E. faecium* E745 [45] in 10-ml BHI supplemented with gentamicin overnight at 37☐, after which 20 μl of the culture was transferred to 10 ml pre-warmed BHI supplemented with 200 μg ml^−1^ gentamicin, which was incubated at 37☐ until the culture reached an OD_600_ of 0.3. After pelleting the cells by centrifugation (3000 g, 5 min), washing once with PBS, and resuspension of the cells in 10 ml PBS, a 3 ml aliquot was transferred to a new tube and incubated at 20☐ for 7 days. The remaining suspension was used for the isolation of genomic DNA. At day 7, the cell suspensions were washed once with PBS and then transferred to 3 ml prewarmed BHI and incubated at 37☐ until OD_600_ 0.3 to recover viable cells. The experiments with the transposon mutant library were performed with four independent replicates.

To prepare libraries for Tn-seq, 2 μg of the extracted genomic DNA was digested for 4 h at 37☐ using 10 U MmeI (New England Biolabs). The DNA was then immediately dephosphorylated by treatment with 1 U calf intestine alkaline phosphatase (Invitrogen) for 30 min at 50☐. DNA was then isolated using phenol-chloroform extraction and subsequently precipitated using ethanol. The DNA pellets were dissolved in 20 μl water. The samples were further processed for Illumina sequencing, including the addition of barcodes, as described previously [45]. Tn-seq libraries were sequenced on one lane of a HiSeq 2500 with 50 nt single-end reads. The sequence reads have been made available on the European Nucleotide Archive with accession number PRJEB37076.

### Tn-seq data analysis

Tn-seq data analysis was performed as described previously [45]. In short, Illumina sequence reads were demultiplexed, based on their barcode, using a customized workflow in Galaxy [52], and 16-nucleotide fragments of each read, corresponding to E745 sequences, were mapped to the E745 genome using Bowtie 2 [53] The results of the alignment were sorted and counted by IGV [54] using a 25-nucleotide window and then summed over the entire gene. Reads mapping to the final 10% of a gene were discarded as these insertions may not inactivate gene function. Read counts per gene were then normalized to the total number of reads that mapped to the genome in each replicate, by calculating the normalized read-count RPKM (Reads Per Kilobase per Million input reads) via the following formula: RPKM = (number of reads mapped to a gene × 10^6^) / (total mapped input reads in the sample × gene length in kbp). Statistical analysis of the RPKM-values between the experimental conditions was performed using Cyber-T [55, 56]. Genes were determined to be significantly contributing to growth in human serum when the Benjamini-Hochberg corrected P-value was <0.05 and the difference in abundance of a gene between day 0 and day 7 was >10 or < −10.

### Construction of targeted deletion mutants

The *usp* deletion mutant was created using CRISPR-Cas9-mediated genome editing as previously described [48]. In short, a CRISPR targeting *usp* was inserted into pVDM1001 by annealing oVDM2001 and oVDM2002 together and ligating this DNA fragment into BsaI-digested pVDM1001 creating pVDM-x*usp*. Next, a DNA fragment (gBlock, Integrated DNA Technologies; Leuven, Belgium) was synthesized containing the 463 bp upstream region of *usp* fused together with the 513 bp downstream region of *usp* **(table S1)**. This DNA fragment was amplified using oVDM2003 and oVDM2004 and was subsequently ligated in SmaI-digested pVDM-x*usp* creating pVDM-*Δusp*. This plasmid was then transformed into *E. faecium* E745 which already carried pVPL3004 [57] with selection for transformants on BHI agar containing 200 μg ml^−1^ spectinomycin and 50 μg ml^−1^ erythromycin at 30☐ for 48h. The deletion of *usp* was confirmed by PCR using oVDM2005 and oVDM2006. The plasmids pVPL3004 and pVDM-*Δusp* were cured from *E. faecium* E745-Δ*usp* by sub-culturing in BHI for 72 hours.

The 848-bp DNA fragment containing the promotor of *usp* and the complete *usp* gene for *in trans* complementation (**Table S1)** was synthesized, and subsequently this fragment was amplified by PCR using oVDM2015 and oVDM2016, and subsequently digested with XmaI and SalI. The digested fragment was ligated into pEF25 to form pEF25-*usp-C*. This vector was subsequently transformed into E745-*Δusp* and transformants were selected on BHI agar supplemented with 200 μg ml^−1^ spectinomycin. The presence of the vector in *E. faecium* E745-Δ*usp* was confirmed by PCR using oVDM2017 and oVDM2018 and the resulting complemented strain was named E745-*Δusp*::*usp-C*.

We were unable to generate an *exuR* deletion mutant with the CRISPR-based genome editing approach described above and we thus used our previously described [49] allelic replacement method, with minor modifications, to generate this mutant. A DNA fragment (Integrated DNA Technologies; Leuven, Belgium) was ordered containing the 496 bp upstream region of *exuR* fused to the 515 bp region downstream of *exuR* **(Table S1)**. The DNA fragment was amplified using oVDM2007 and oVDM2008, digested with XhoI and XmaI and subsequently ligated into similarly digested pWS3 to create pVDM-*exuR*. The gentamicin resistance cassette from pGPA1 was amplified using oVDM2009 and oVDM2010, followed by digestion with EcoRI. This DNA fragment was then ligated into similarly digested pVDM-*exuR*, resulting in pVDM-*exuR+G*. E745 was transformed with pVDM-*exuR+G* and transformants were selected at 30☐ on BHI supplemented with 200 μg ml^−1^ spectinomycin. The presence of pVDM-*exuR+G* in the transformants was confirmed by PCR using oVDM2011 and oVDM2012. To induce a double crossover event to replace *exuR* with the gentamicin resistance cassette, a colony harbouring pVDM-*exuR+G* was used to inoculate 200 ml BHI medium and then subcultured for 72 hours at 37☐ after which the culture was plated on BHI with 300 μg ml^−1^ gentamicin. At least 100 colonies were transferred to BHI agar containing either 300 μg ml^−1^ gentamicin or 200 μg ml^−1^ spectinomycin. Colonies that were resistant to gentamicin but not spectinomycin were checked for successful double crossover events using oVDM2013 and oVDM2014. The resulting mutant was named E745::*ΔexuR+G* and was consequently transformed with pCRE-*lox* to remove the gentamicin cassette from the genome, as described previously [49]. Removal of the gentamicin cassette was confirmed using PCR using oVDM2013 and oVDM2014 and curing of pCRE-*lox* after removal of the gentamicin cassette was performed as described previously [49].

### Presence and absence of genes involved in survival during starvation in *E. faecium* genomes

Abricate (https://github.com/tseemann/abricate, version 0.8) was used to identify presence of the 18 *E. faecium* E745 genes that had the largest role in survival in PBS, as determined by Tn-seq, against the draft assemblies from 1,644 *E. faecium* genomes that represent the global diversity of the species *E. faecium* [58]. We considered a minimum identity and coverage of 95% and 80%, respectively, to consider a gene as present in a particular draft assembly. To visualize the presence and absence of these genes in the context of the phylogeny of *E. faecium*, we used the neighbour-joining tree described in [58], based on a RAxML tree-based of 955 *E. faecium* core genes, and used the R package ggtree (version 1.14.6) to plot the presence of the different E745 genes.

## Notes

### Competing Interest Statement

The authors have declared no competing interest.

