## Supplementary material for "Conditionally essential genes for survival during starvation in *Enterococcus faecium* E745": Figure S1

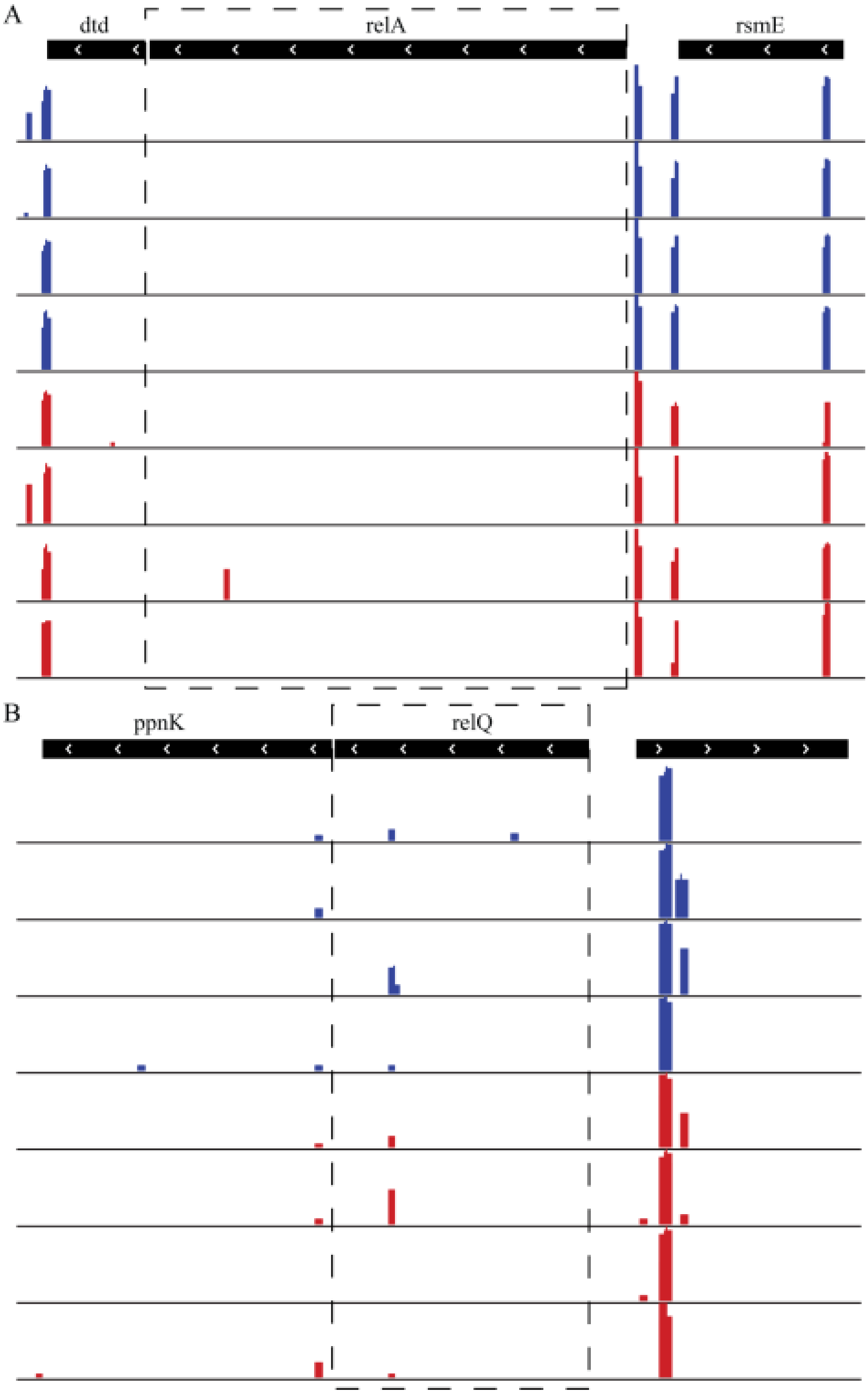


**Figure S1: Overview of transposon insertions in the genetic region of *relA* (EfmE745_02431) and *relQ* (EfmE745_02408).** Transposon abundance for *relA* is shown in panel A and for *relQ* are shown in panel B. Black bars with arrows indicate genes and their direction of transcription. Underneath the genes is a representation of the relative abundance of transposons on a $\log_{10}$ scale. Blue and red bars denote the abundance of transposon insertion mutants at day 0 and after starvation for 7 days, respectively.
